# Gram-positive bacteria are primed for surviving lethal doses of antibiotics and chemical stress

**DOI:** 10.1101/2024.05.28.596288

**Authors:** Manisha Guha, Abhyudai Singh, Nicholas C. Butzin

**Affiliations:** Department of Biology and Microbiology; South Dakota State University; Brookings, SD, 57006; USA; Electrical & Computer Engineering; University of Delaware; Newark, DE 19716; USA; Department of Chemistry and Biochemistry; South Dakota State University; Brookings, SD, 57006; USA

## Abstract

Antibiotic resistance kills millions worldwide yearly. However, a major contributor to recurrent infections lies in a small fraction of bacterial cells, known as persisters. These cells are not inherently antibiotic-resistant, yet they lead to increased antibiotic usage, raising the risk of developing resistant progenies. In a bacterial population, individual cells exhibit considerable fluctuations in their gene expression levels despite being cultivated under identical, stable conditions. This variability in cell-to-cell characteristics (phenotypic diversity) within an isogenic population enables persister cells to withstand antibiotic exposure by entering a non-dividing state. We recently showed the existence of “primed cells” in *E. coli*. Primed cells are dividing cells prepared for antibiotic stress before encountering it and are more prone to form persisters. They also pass their “prepared state” down for several generations through epigenetic memory. Here, we show that primed cells are common among distant bacterial lineages, allowing for survival against antibiotics and other chemical stress, and form in different growth phases. They are also responsible for increased persister levels in transition and stationary phases compared to the log phase. We tested and showed that the Gram-positive bacterium *Bacillus megaterium*, evolutionarily very distant from E. coli, forms primed cells and has a transient epigenetic memory that is maintained for 7 generations or more. We showed this using ciprofloxacin and the non-antibiotic chemical stress fluoride. It is well established that persister levels are higher in the stationary phase than in the log phase, and B. megaterium persisters levels are nearly identical from the early to late-log phase but are ∼2-fold and ∼4-fold higher in the transition and stationary phase, respectively. It was previously proposed that there are two distinct types of persisters: Type II forms in the log phase, while Type I forms in the stationary phase. However, we show that primed cells lead to increased persisters in the transition and stationary phase and found no evidence of Type I or II persisters with distant phenotypes. Overall, we have provided substantial evidence of the importance of primed cells and their transitory epigenetic memories to surviving stress.

## Introduction

Antibiotics combat bacterial infections in animals and humans either by eradicating the bacteria or impeding their ability to thrive and reproduce. Bacterial cells employ various mechanisms to defend themselves against antibiotic attacks, the two most common ways being resistance and tolerance. Some bacteria are naturally resistant, but others develop antimicrobial resistance by acquiring resistant genes[1-5], or through the accumulation of mutation(s) in the bacterial genome, which allows bacteria to counter the antimicrobial drugs[3-5]. According to the Centers for Disease Control and Prevention (CDC), unfortunately, over 2.8 million antimicrobial-resistant infections occur annually in the US, claiming more than 35,000 lives. These numbers are likely extremely underestimating the problem because not all individuals who were infected are tested for antibiotic-resistant bacteria. Recent studies show that worldwide antibiotic-resistant bacteria cause more fatalities compared to other infectious diseases like AIDS, with global death projections to be at 10 million per year by 2050[6-8]. It is important not to underestimate how much of a growing threat this is. For example, cancer is currently the 2nd leading cause of death worldwide, killing about 9.5 million people each year[9, 10]. The antibiotic threat is posed to kill just as many people in 25 years unless we develop new tools to kill these pathogens. We need to understand better the mechanism that allows bacteria to survive antibiotics to build these new tools to combat this growing threat.

Here, we focus our efforts specifically on antibiotic persisters because they are a major driver of antibiotic resistance development and because persisters have been shown to be the cause of recurrent bacterial infections[11-13]. The addition of lethal antibiotics dosage results in a biphasic death curve: Phase 1 contains non-persister cells (susceptible cells and short-term tolerant cells) that are quickly killed off. Phase 2 contains persister cells that take an exceptionally long time to be killed compared to their susceptible counterparts[14, 15]. Cells in Phase 1 and Phase 2 are genetically identical cells (Fig. 1A; Fig. S2D). Though persisters are nonresistant to the antibiotic and are genetically identical to the antibiotic-susceptible cells, incomplete treatment (use of too low of an antibiotic concentration or not treating long enough to kill all persisters) can result in persister cell survival. These cells emerge after antibiotic treatment, and their offspring have a higher mutation rate and are more likely to mutate or accumulate resistance genes from their microenvironment[16-18].

**Fig. 1.**
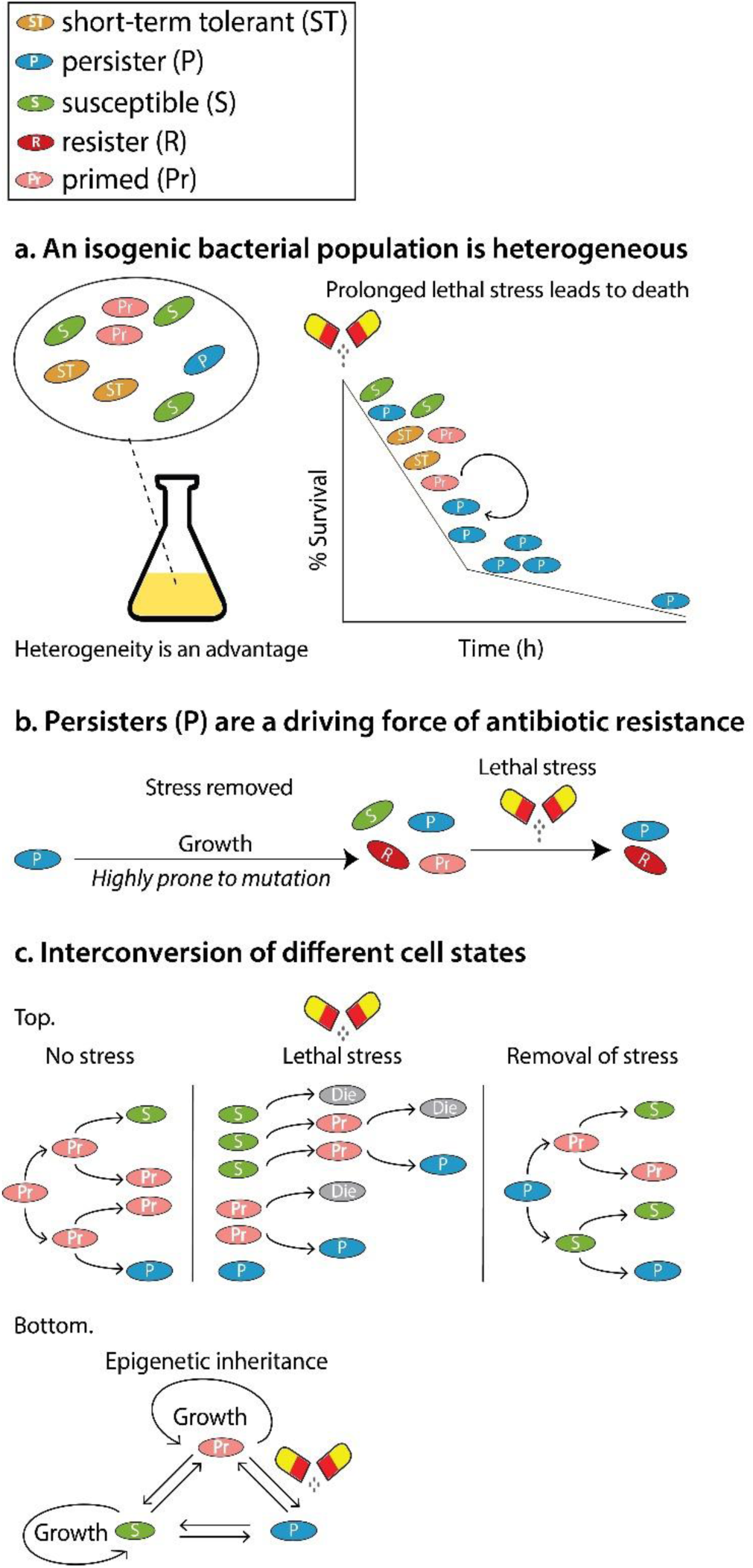
**(A). An isogenic bacterial population is heterogeneous.** A bacterial population consists of antibiotic-susceptible (S), short-term tolerant (ST), primed (Pr), and persister (P) cells (sometimes resistant (R) cells as well). A bacterial population shows a typical biphasic death curve upon the addition of lethal doses of antibiotics. The susceptible cells die out the fastest, followed by short-term tolerant cells (phase 1 of the death curve). The persisters (both initially existing as well as converted from the primed cells) survive much longer than other cells (phase 2 of the death curve). **(B). Persisters (P) are a driving force of antibiotic resistance.** Upon stress, only the persisters (and resisters) survive. With prolonged stress, the persisters eventually die out, but the resisters survive. Upon removal of stress, persisters revert to normal cells, divide, and are more likely to accumulate mutations, forming resistant cells. **(C). Interconversion of different cell states. Top:** Primed cells are dividing cells that are prepared to combat stress before encountering it. They can give rise to progeny of primed cells, persister cells, or susceptible cells, but this may depend on the environment. We do not know all the states cells can transition to and from, but here we show several possibilities. **Bottom:** Due to phenotypic heterogeneity, bacterial cells exist in interchangeable states. We have demonstrated that primed cells can give rise to more primed cells through epigenetic inheritance and that primed cells can be converted to persisters and susceptible cells. We have shown that susceptible cells are converted to primed cells, but we do not know if persisters are directly converted to suspensible cells or if they are first converted to primed cells. We also do not know if persister cells are converted to primed cells directly.

For the longest time, it was assumed that clonal bacterial cultures would behave in an identical manner; however, there is a growing realization of the importance of heterogeneity in an isogenic bacterial population. In general, even pure cultures of bacteria in the exponential (Log) phase of growth are heterogeneous in nature (Fig. 1A). This is true both under well-defined laboratory conditions as well as in natural environments[19, 20]. Cells in a bacterial population exhibit variability in gene expression (either at the transcription or translation level), leading to phenotypic heterogeneity[21]. If this noise in gene expression is amplified through positive feedback, then a graded response shifts to a binary response, leading to the establishment of a dynamic bistability wherein a cell can stay in two different interchangeable states[22-24] (Fig. 1C). Heterogeneity leads to reduced fitness in an optimal, steady-state environment, but increases chances of survival during environmental disturbances, a phenomenon called bet-hedging[25-27]. Bacterial persistence is a classic paradigm of bet-hedging strategy where a small fraction of cells in a growing culture under normal conditions is in a state of arrested division with differential gene expression[28, 29], or a small fraction of cells are primed to enter the persister state[30]. In the presence of antibiotics, these physiological conditions aid persisters in outliving susceptible cells. Once the stress is removed, persisters can switch to a dividing state (non-persister state), forming a population of antibiotic-susceptible cells and persisters[31, 32] (Fig. 1B-C). This re-establishes bacterial infection after treatment, leading to the prolonged usage of antibiotics.

Phenotypic variability has been reported to be both a chance event due to stochastic fluctuations in gene expression and an evolvable trait[33], in different contexts[22, 34]. It has been 80 years since persistence was first demonstrated in 1944[35], but the mechanism of persister formation is still being debated. We recently demonstrated that some cells, called “primed cells”, are prepared for stress before the stressor hits by using the model Gram-negative bacteria *E. coli*[30]. We wanted to know if primed cells are unique to *E. coli* and unique to antibiotics. In this work, we tested primed cell formation in *Bacillus megaterium* (also known as *Priestia megaterium*) because it is a very distantly related organism to *E. coli*. *E. coli* is gram-negative, while *B. megaterium* is gram-positive.

Progress in single-cell technologies has revealed notable variations in phenotype and expression patterns among individual cells within a given isogenic cell population[36-42]. This method has recently been used to explore cancer persistence[43-46] (cancer persistence is different from bacterial persistence, but they are both examples of a small fraction of a population of cells with the ability to survive lethal stress). However, single-cell sequencing methods can provide only a static impression of different cell states[47]. We leveraged a modified Luria-Delbrück Fluctuation Test (FT) to characterize the dynamics of intercellular transitioning from one cell state to another over multiple generations on a temporal scale. The classical FT devised by Luria and Delbrück demonstrated that genetic mutations for survival occur randomly and not in response to the selection pressure[48]. At this time, there was a debate between two conflicting theories: one stated that resistance mutations occurred randomly before the selection pressure (Darwinian theory), and the other stated that they were induced by the selection pressure (Lamarckian theory). Single *E. coli* cells were grown into clones and infected by T1 phage (viruses that infect and kill *E. coli* cells). If each bacterial cell has a small and independent chance of acquiring phage-induced mutation, then the count of resistant bacteria in each clonal population should adhere to Poisson distribution. On the other hand, if mutations occur randomly, the count of mutant cells will vary significantly among clonal populations based on when the mutation emerged during colony amplification. Their experiments yielded a skewed non-Poissonian distribution in the number of mutant cells, confirming the Darwinian theory of evolution (Fig. 2A). The FT showed the potential of fluctuation-based analyses to unveil random processes that are latent. Clone-to-clone fluctuations can also be used to calculate the rate of switching between the two cellular states[49]. While Luria and Delbrück worked on irreversible, genetic changes causing phage resistance, we employed a modified FT to determine the transitory presence of “primed cells” (a cell state arising from reversible, non-genetic processes) in a bacterial population before antibiotic treatment (Fig. 2B). Primed cells are a subpopulation of bacteria priorly prepared to combat stress, and primed cells give rise to persisters[30].

**Fig. 2.**
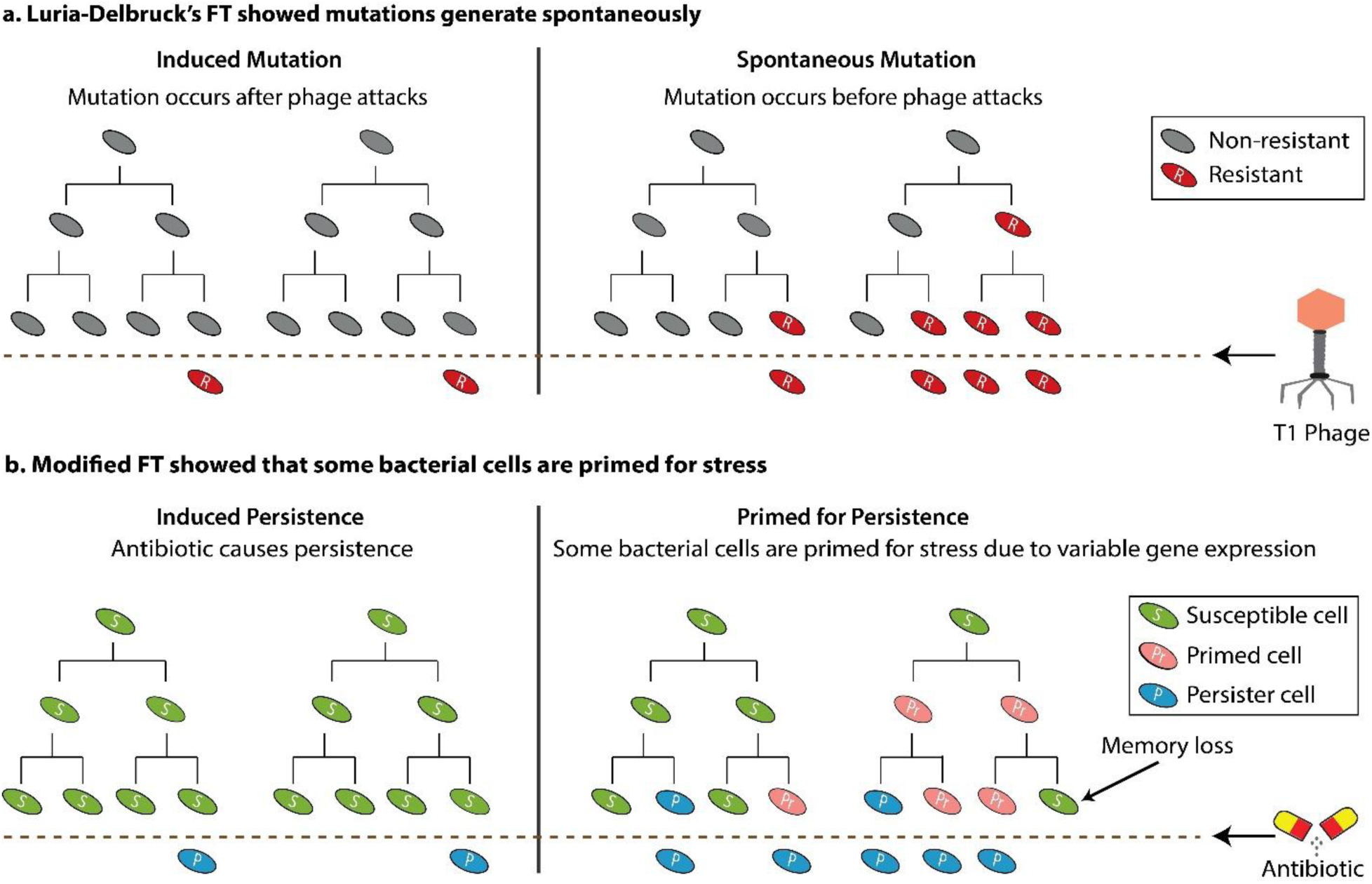
**(A). Luria-Delbrück’s FT showed mutations generate spontaneously.** The classic Luria-Delbrück fluctuation test (FT) is where each clone starts from a single *E. coli* cell and is infected by T1 phage. If resistance mutations are virus-induced, the number of resistant cells would follow a Poisson distribution across clones. Alternatively, if mutant cells arise spontaneously before viral exposure, there will be considerable clone-to-clone fluctuations in the number of resistant cells. Their results showed that spontaneous mutation theory is correct in this case. **(B). Modified FT showed that some bacterial cells are primed for stress.** FT can determine if some cells are primed for stress. This design has a similar experimental setup as a. but measures variation in persister cells across the clones due to antibiotic stress. If persisters are antibiotic-induced, then the number of persisters would follow a Poisson distribution across clones. Alternatively, if there are primed cells that allow them to prepare for stress before antibiotic exposure, there will be considerable clone-to-clone fluctuations in the number of persisters. Our results show the presence of primed cells.

In this study, we utilized a modified FT to determine the presence of primed cells in *Bacillus megaterium* culture at distinct phases of bacterial growth, namely log phase, late-log to early-stationary transition phase (henceforth referred to as the transition phase), and stationary phase. We show that primed cells have a transient memory and are effective in combating both antibiotic and chemical stress (fluoride, F^-^). In the post-log phase of growth, the number of primed cells increases proportionally with the number of persisters. Combining the FT results with mathematical support, we have elucidated how stochastic, heritable, transient phenotype switching leads to variation in persister levels on exposure to antibiotics. Understanding this inherent survival strategy can delineate stress response mechanisms in bacteria, which is crucial for drug development against persisters and effective treatment of chronic recurrent infections[50-58].

## Results and Discussion

### B. megaterium culture consists of “primed cells” prepared for lethal antibiotic stress prior to treatment

We previously showed that primed cells allow *E. coli* to prepare for antibiotic stress[30], so we used a similar procedure to test if *B. megaterium* also forms primed cells. This involves a comparison of the persisters formed from the Noise Control (NC) to those formed from a modified Fluctuation Test (FT; single cell-level persister percentage variation assay) (Fig. 3A-B). On average, a bacterial culture usually has a narrow range and fold difference between the maximum and the minimum persister percentages obtained from that population, and this range is called the Noise Control (NC). A *B. megaterium* culture was grown to mid-log phase (OD_600_ of about 0.4-0.5) and then distributed into 48 wells of a 96-well optical bottom plate. The wells containing the bacterial culture were then treated with ciprofloxacin (Cip) for 3 h to kill non-persister cells. This population-level persister percentage variation assay resulted in an NC of ∼1.8-fold.

**Fig. 3.**
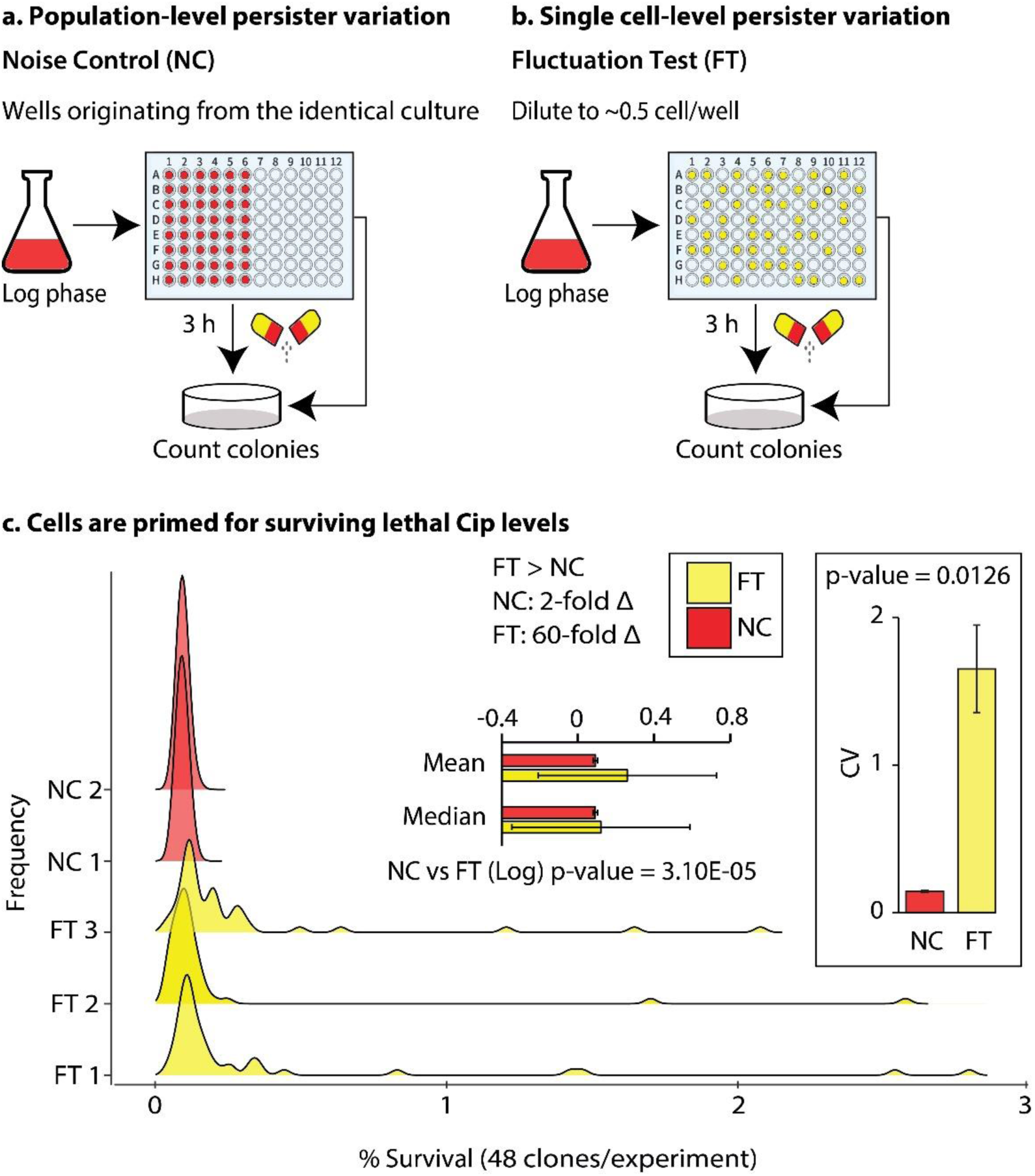
**(A). Population-level persister variation.** Noise Control (NC): population-level persister percentage variation in a bacterial population. **(B). Single cell-level persister variation.** Fluctuation Test (FT): single cell-level persister percentage variation in clonal populations. **(C). Cells are primed for surviving lethal Cip levels.** Some *B. megaterium* cells are primed (prepared) before Cip treatment; NC: ∼2-fold, FT: ∼60-fold. The mean, standard deviation, and CVs were obtained through bootstrapping with a 95% confidence interval. A t-test between the variances of NCs and FTs showed that they are significantly different (p-value<0.05).

The FT involves diluting a bacterial culture down to a single cell in a 96-well optical bottom plate. To do this, we dilute cells to ∼0.5 cells per well. Cells cannot be split in half and remain viable; so on average, there will be about 1 cell per 2 wells[59, 60] and result in ∼48 out of 96 wells having growth. The clonal cultures were then grown to an OD_600_ of about 0.4-0.5 and then treated with Cip for 3 h to perform a persister assay. Because not all wells reach the same cell density simultaneously, two 96-wells were used for this assay, but only 48 wells with OD 0.4-0.5 were treated with Cip.

The difference between the FT and NC is striking. The FT results for *B. megaterium* with Cip resulted in a ∼60-fold variation in persistence (Fig. 3C) compared to the ∼1.8-fold variation with NC. This was tested multiple times, and the results were consistent. Since the bacterial culture was diluted down to single cells and subcultured to the log (exponential) phase, any residuals from the stationary phase were lost. The *B. megaterium* results are consistent with *E. coli* with ∼80- and ∼60-fold differences in persister percentages with ampicillin and apramycin, respectively[30].

Two different hypotheses explain the high variation in the FT. First, though unlikely, the high fluctuations in a few clonal populations were perhaps due to some mutation leading to resistance rather than persistence. Second, some bacterial cells in a population were already prepared or primed for combating lethal doses of antibiotics. If the cells from the high persister culture were resistant, then their Minimum Inhibitory Concentration (MIC) are highly likely to increased[61]. The MIC of Cip for a high persister *B. megaterium* clonal culture was tested and found to be identical to that of susceptible *B. megaterium* culture (Fig. S1A), thus mutation is very unlikely. Multiple colonies from high-persister plates were picked and streaked on Cip plates. No growth was observed (Fig. S1B). An FT was performed with a high-persister clonal culture. If mutations were the cause, then the average persister percentage of the high-persister clone should increase, and the range of persister percentage should decrease compared to the wild-type clonal populations. However, when repeating the persister assay with the high-persister clone, there was no apparent difference between it and wild-type (Fig. S1C). These experiments ruled out the first hypothesis, thereby demonstrating that some cells in a bacterial population are priorly prepared for antibiotic stress. These results match our previous work with *E. coli* primed cells[30], so we called these stress-prepared cells *Bacillus* primed cells.

In our previous work with *E. coli* primed cells, the antibiotics ampicillin (β-lactam class) and apramycin (aminoglycoside) were used, which target peptidoglycan synthesis and 30S ribosomal subunit respectively[30, 62, 63]. Here, we used Cip, a quinolone that inhibits DNA gyrase[64], to identify Gram-positive primed cells, which further supports that cells prepare prior to a generalize antibiotic stress, and not to any particular class of antibiotic.

### Primed cells have short-term memory

We previously demonstrated that primed cell memory of persistence was passed down to the progeny of *E. coli* cells, and the high variation range in persistence was not random in every generation[30]. We wanted to know if the same was true for *B. megaterium*. Our null hypothesis was that there is no memory, and the variation range in persistence is random. We used a method that allows us to compare persister levels between replicas to test our hypothesis (Fig. 4A). A *B. megaterium* culture was diluted to single cells and grown to mid-log phase (OD_600_ of about 0.4-0.5). It was then divided into two 96-well microplates, diluted (to 1:2, 1:10, or 1:100), and allowed to grow. Persister assays were performed with both the replicates using Cip; however, each separate plate (Replica Plate 1 and Replica Plate 2) was allowed to grow independently from each other. If there was a memory in the system, the corresponding well in Replica Plate 1 should have similar levels of persisters as each well in Replica Plate 2. However, if there was no memory in the system, the corresponding well in Replica Plate 1 should not have similar levels of persisters as each well in Replica Plate 2. We can test for memory by seeing if there is a correlation between the persister levels in Replica Plates 1 and 2.

**Fig. 4.**
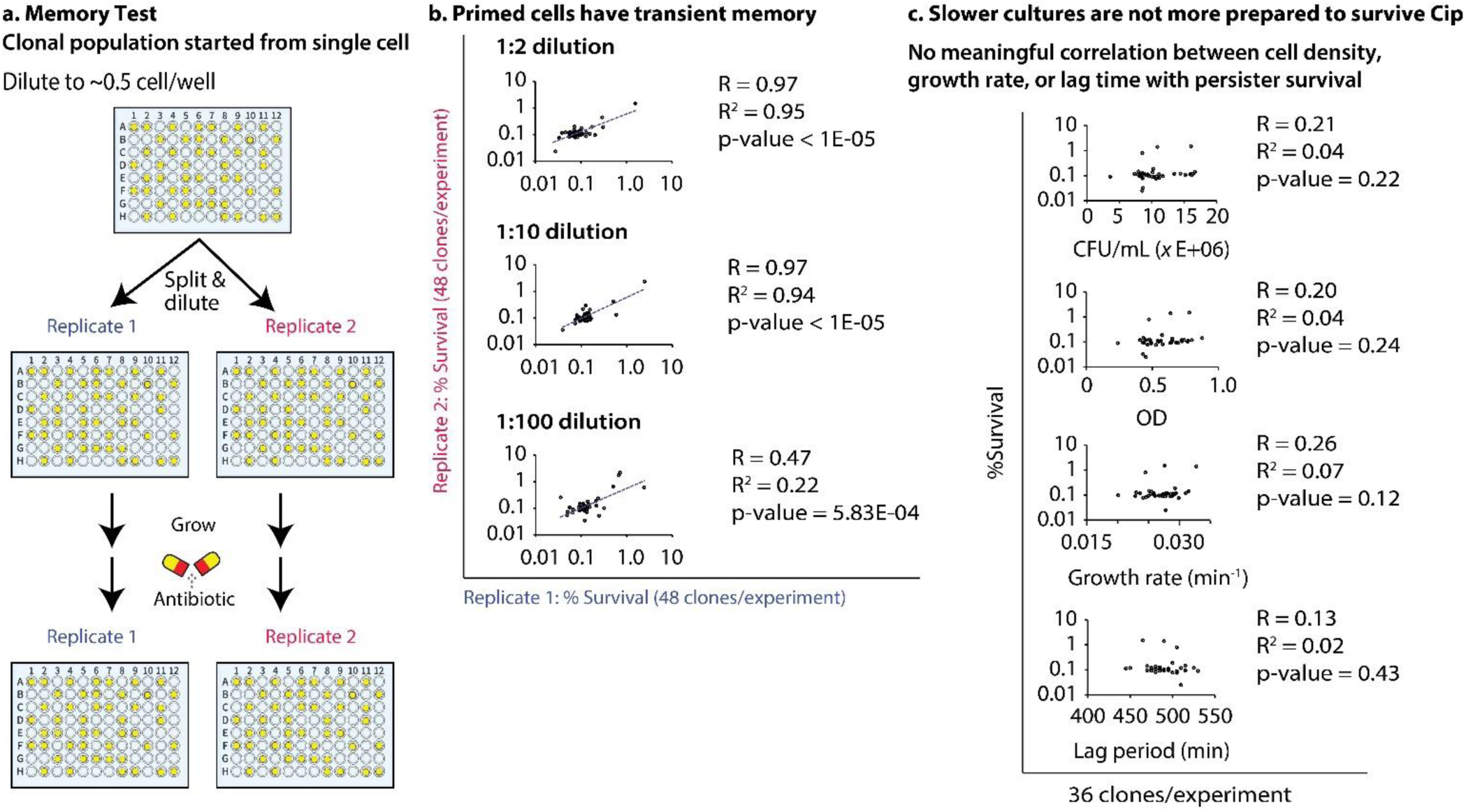
**(A). Memory Test.** Single cell-level persister percentage variation in clonal populations of two replicates. **(B). Primed cells have transient memory.** Primed cells of *B. megaterium* have transient memory before antibiotic addition, which gets passed for >7 generations. The T-test between the FTs of two replicates for each dilution showed that they were not significantly different. Both the x and y axes are log_10_ scaled. **(C). Slower cultures are not more prepared to survive Cip.** *B. megaterium* cultures with different cell densities (as indicated by CFU/mL as well as OD), grown over the same duration, did not show any correlation with persister percentages. The growth rate of the individual cultures, as well as their lag period, also did not show any correlation with persister percentages. The y-axis is a log_10_ scale.

The persister levels in both the replicates strongly correlated when cells were split and diluted 1:2 and 1:10, defying our null hypothesis. (Fig. 4B). The correlation was less when diluted to 1:100, but this was not surprising because the memory is transitory. These results also support our previous results that mutations are not the cause of high variation in persister percentages in clonal populations because even with 1:100 dilution, the clones did not show any resistant phenotype. Our *Bacillus* work is similar to our previous work with treating *E. coli* with ampicillin and apramycin, where primed cell memory was lost before 1:100 dilution[30]. Since persisters are genetically identical to the antibiotic susceptible cells, the logical conclusion is that the memory is epigenetic. Epigenetic memory is a short-term behavior change that propagates this transient and stable cellular phenotype from parent to daughter cells[65]. One primed cell will divide to become two primed cells, and so on, but eventually, that memory that resulted from non-DNA change is lost. In this case, the memory is maintained for 7 generations or more.

There are a lot of possible ways through which epigenetic inheritance can take place in bacteria[66-69]. Multisite phosphorylation[70, 71] and ploidy[72] have been implicated in the regulation of phenotypic heterogeneity in some cases, but there are other options. This inheritable transient state may be favored in evolution as a survival strategy compared to DNA mutation, as it does not necessitate a long-term commitment[73]. In this work, we did not explore the exact epigenetic inheritance mechanism in play during primed cell division.

### Primed cell numbers increase in post-log phase, resulting in more persisters in the transition and stationary phases than in the log phase

Bacterial persistence levels are known to increase in the late-exponential phase in some bacteria[32, 74, 75], and *Pseudomonas aeruginosa* may modulate persister frequency partly via quorum sensing[76, 77]. These are density-dependent responses. We wanted to know if *B. megaterium* persister levels and primed cell number varied based on cell density. We see no difference in persister percentages with OD_600_ 0.25-0.8 (log phase) in the presence of Cip (Fig. 5B). This was not particularly surprising because in the log phase, in a lab setting, the environmental condition is kept constant and nutrient levels are plentiful. However, we were more interested in what would happen to the persister and primed cell numbers when cells were in a more stressful environmental condition such as transition and stationary phases (Fig. 5A).

**Fig. 5.**
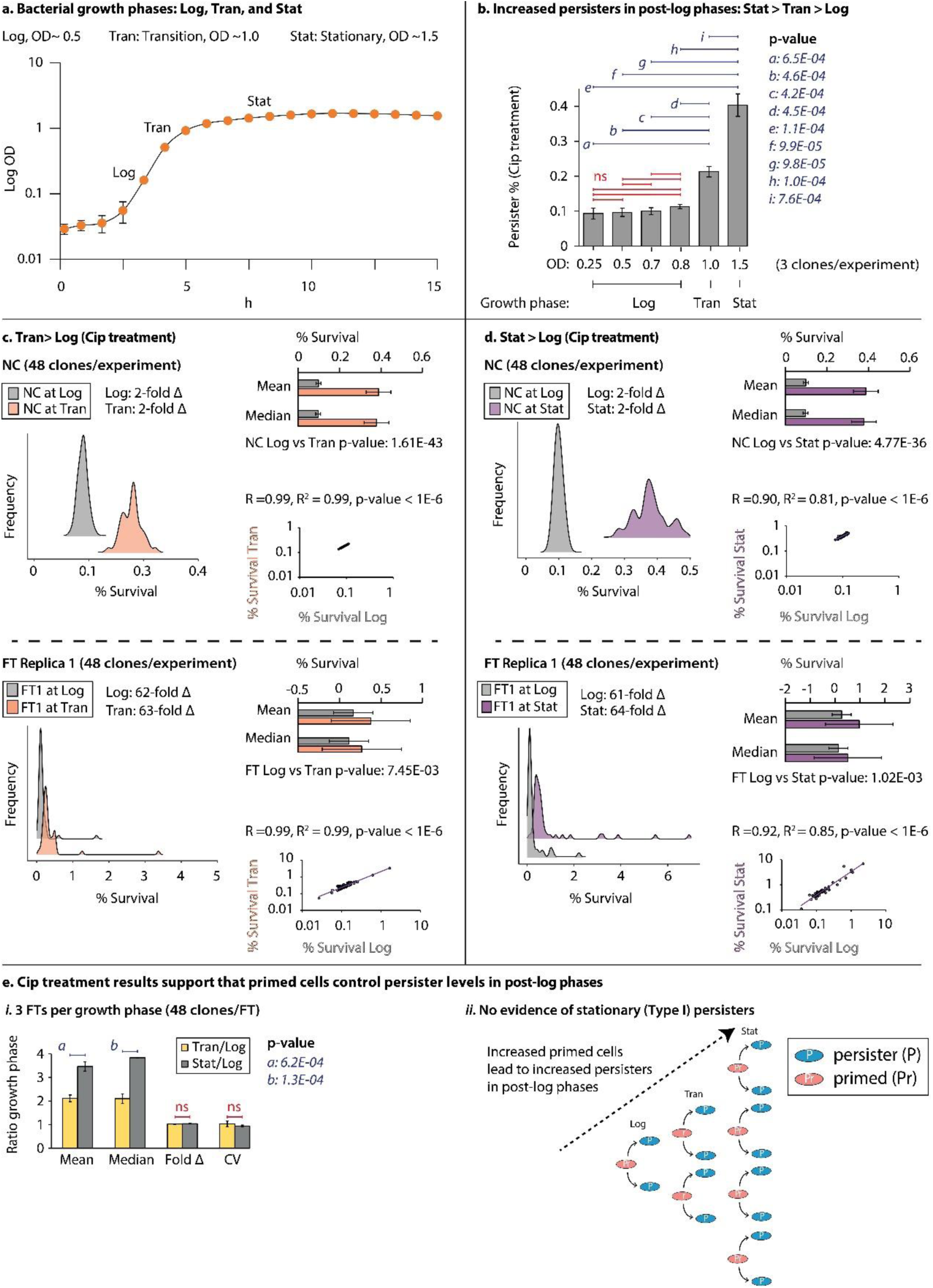
**(A). Bacterial growth phases: Log, Tran, and Stat.** *B. megaterium* growth curve with 3 different growth phases: Log, Transition (Tran), and Stationary (Stat). **(B). Increased persisters in post-log phases: Stat>Tran>Log.** The persister numbers remained constant during the log phase of growth and increased thereafter (n=3; the non-significant (ns) p-values are in Fig. S2B). **(C, D). Tran>Log and Stat>Log (Cip treatment).** Results of population-level (NC) and single cell-level (FT1) persister percentage variation assays at Log vs Tran, and Log vs Stat phases of growth with Cip. Compared to the log phase, the average persister percentage in the Tran and Stat phases are ∼2-fold and ∼4-fold higher, respectively. However, the fold difference between the highest and lowest persister percentages (fold Δ = Max. persister % / Min. persister %) remains the same in all the phases. There is a proportional shift in the average persister percentage along with the number of primed cells in the post-log phases. The persister percentages of Log vs Tran or Stat correlate strongly as indicated by R square and p-values. Both the x- and y-axes are log_10_ scaled. Two more replicates (FT2, FT3) for C. and D. are in Fig. S3A-B. **(E). Cip treatment results support that primed cells control persister levels in post-log phases. (i).** The mean, median, fold change (Fold Δ) of persister % and CVs of Tran/Log and Stat/Log phases were calculated through bootstrapping with a 95% confidence interval. The mean and median persister % increase post-log phase, but the fold Δ and CVs remain constant. The p-values labeled ns for the comparison between Tran/Log and Stat/Log are 0.25 for Fold Δ and 0.29 for CV. **(ii).** From the Log to Tran phase to Stat phase, the numbers of primed cells proportionally increase with the number of persisters.

The average persister percentage of *B. megaterium* culture increased ∼2-fold in the transition phase (OD_600_ ∼1.0) and ∼4-fold in the stationary phase (OD_600_ ∼1.5) as compared to that in the log phase (Fig. 5C-D). The transition and early stationary phases were chosen to avoid sporulation (Fig. S2. iii) and yet are sufficiently more heterogeneous than the log phase. Despite the average persister percentage increasing post-log phase, for both NC and FT, the CVs and the fold change (the differences between the highest and lowest persister percentages) remained similar to that in the log phase (Fig. 5E. i). Only the mean and median were different.

The overall results show that primed cell number increases in later phases. This increased primed cell number leads to more persisters (Fig. 5E. ii). It is often cited that there are two types of persisters: Type I persisters are non-growing cells that form in the stationary phase, while Type II persisters originate from slow-growing cells that generate continuously[31]. Our results show that there are more persisters in the stationary phase but provide evidence that persisters in stationary and log phases are not different types (Type I or II) based on 3 points:

1. The primed cell numbers control the total number of persisters formed regardless of the phase of growth they are in. If Type I and Type II persisters are distinct in their formation, then how can the primed cell numbers control both?
2. Overall, primed cells are not slow growing in the log phase but are the primary factor in controlling the persister levels. This is also supported by our previous results with *E. coli,* where antibiotics ampicillin (β-lactam class) and apramycin (aminoglycoside) lead to massive changes in cell survival numbers[30].
3. The “macro” growth rate does not appear to affect the primed cell or persister level. It is important to note that the “macro” growth rate is not a reflection of the individual cell rate. Thus, our results do not rule out the possibility that individual cell growth rates could be related to persister cell formation. However, these results do demonstrate that primed cell numbers are more substantial in driving cell survival than slower growth rates.

### Persister numbers do not always depend on the time of growth

In all previous experiments, the cell density was kept very similar (∼ODs around 0.5) to minimize potential variations between wells. When a *B. megaterium* culture is diluted to single cells in a 96-wells microplate (∼0.5 cells/well) and incubated, they do not all grow at a similar rate and thus reach higher cell densities at different times. We were interested in whether the overall density or growth of a log culture affected primed cells or persister cell numbers. So, the FT was performed with 36 clonal populations grown for the same time period but had OD_600_ ranging from 0.3 to 0.8 (Fig. S2A). There is no correlation between cell density (based on OD or CFU/ml) and persister level (Fig. 4C).

According to previously well-cited literature, slower growth rates lead to higher persisters[78-80]. Therefore, clonal populations with lower values of OD_600_ should have higher persister percentages. In addition, the goal is to have 1 cell/well, but this will not always be the case. A well may reach a higher density than another because it contains more than one cell, but it does not grow faster. A highly effective way to control more than 1 cell/well and to determine if slow growth leads to more persisters is to compare growth rates. The growth rate of individual wells did not correlate with the persister numbers (Fig. 4C). This is in direct contrast to literature claiming that a slow growth rate leads to greater persistence. Another possibility is that the delay time before cells divide leads to higher persisters. This delay can be measured by determining the lag time. The lag period of individual wells did not correlate with the persister numbers (Fig. 4C).

These FT results show that slow growth does not lead to appreciable changes in persister or primed cell levels. This type of experiment has a limitation: we do not visualize the growing rate of single cells. However, new research has shown that antibiotic lethality is more depedent on the metabolic state than growth rate [81], which is consistent with our results. Based on our results, we concluded that the majority of persisters being formed are not slow-growing, and if slow-growing cells lead to higher persister levels, this is only a minor fraction of the overall number of persisters being formed.

### Primed cells are prepared for stress in general

All the experiments with *B. megaterium* and our previous work with *E. coli[30]* focused on antibiotic survival. Persistence may be a general survival strategy, and previous researchers have identified persisters to non-antibiotic stresses[82]. We wanted to know whether primed cells of a bacterial population are prepared for survival against other stressors, so we tested lethal concentrations of fluoride. Fluoride is an agent commonly found in toothpaste and water supply and is used to prevent dental caries in humans[33, 83-87]. Fluoride in large quantities is bactericidal, mainly by reducing the acid tolerance in bacteria through a combination of enzyme inhibition, altering membrane permeability and inhibition of certain nutrient metabolism[88-99].The fold differences between the NC and FT were ∼2.6-fold and ∼78-fold, respectively (Fig. 6A). This demonstrated that primed cells are prepared, in general, for many kinds of stressors. Maintaining elevated levels of primed cells provides selective advantage to a bacterial population by imparting phenotypic plasticity in the face of environmental stress.

**Fig. 6.**
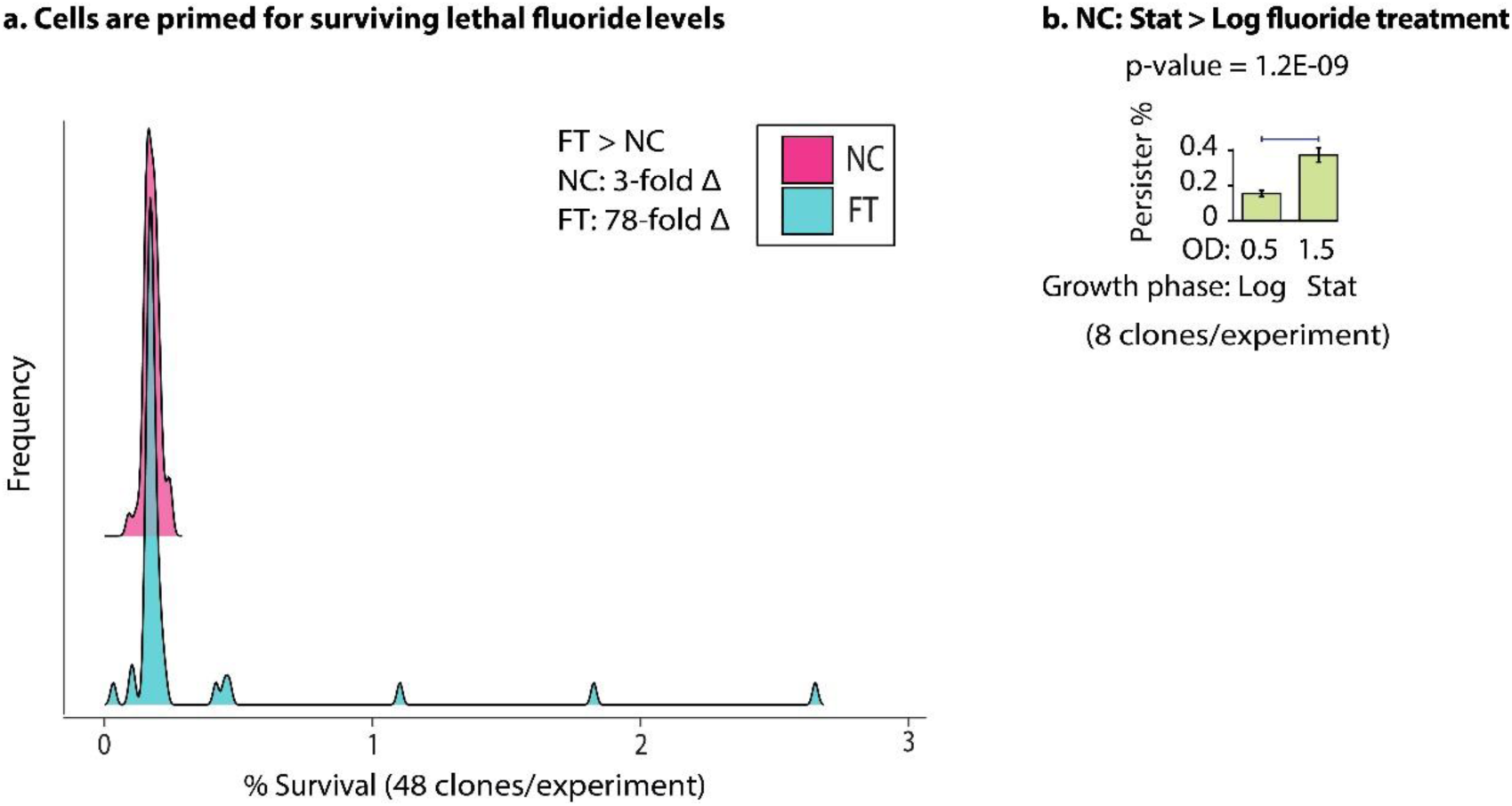
**(A). Cells are primed for surviving lethal fluoride levels.** Some *B. megaterium* cells are primed (prepared) before fluoride treatment (a non-antibiotic chemical stressor); NC: ∼3-fold, FT: ∼78-fold. The differences in the mean and median of NC and FT are represented in bars. **(B). NC: Stat>Log fluoride treatment.** The persister percentages of B. megaterium increase when treated with fluoride from log (OD_600 nm_ 0.5) and to stationary phase (OD_600 nm_ 1.5) by ∼ 2.5-fold (n = 8).

We have demonstrated that the primed levels lead to increased persisters in the stationary phase compared to the log phase when using the antibiotic Cip. This increase could be a general phenomenon or unique to antibiotics, so we tested cells with lethal fluoride. The number of primed cells in the stationary phase was higher with fluoride than in the log phase, which shows that this is not an antibiotic-specific phenomenon (Fig. 6B).

## Outlook

Bacterial populations employ multiple approaches to survive stressful environmental conditions. One of them is bet-hedging, where a bacterial population has a variant phenotype that is maladjusted for the current environment but suited for a changed environment. This phenotypic plasticity is a result of noise in gene expression[21, 25, 100, 101]. At first glance, noise might not seem like a driver of such a crucial phenotypic heterogeneity that allows some cells (persisters) to survive lethal doses of antibiotics. Although, in general, certain aspects of this variability can be associated with external factors such as cell size, cell-division phase, and the extracellular microenvironment, an increasing body of evidence indicates that stochastic processes inherent in gene expression play a significant role in generating random fluctuations (noise) in the levels of gene products[73, 102-116]. Understanding the elemental mechanisms of heterogeneity is pivotal for developing drugs against persistence.

A simplified way to think about this is to imagine the expression of 1 gene in cells with identical genomes and in the same environment at the same time. In the Central Dogma of Life, DNA is replicated to DNA by DNA polymerase, DNA is transcribed to RNA by RNA polymerase, and RNA is translated into protein by ribosomes. It would be ridiculous to think cells have identical levels of these components with identical activity. At each of these steps, there is variation: variation in the amount of DNA, the DNA polymerase activity, the RNA polymerase activity, the ribosomal activity, and the multitudinous of other molecules that come into play. At each step in the process, from DNA to protein, there are variations, and these variations add up. The result is that, on average, identical cells have nearly identical behavior, but within a large population, there will be outliers. These outliers lead to primed cells and persisters. In this example, we focused on the variation in a single gene, but this is oversimplified. Cells have many genes and contain many molecules that vary slightly in individual cells in the same environment. Cells experience slight variations in their external and internal environment, which means that there are many opportunities for these outliers to form in a large population. What we have shown here is that these outliers (primed cells) not only form due to variations but can pass their uniqueness (behavior) down for several generations, and that epigenetic memory allows these cells to be more prepared to survive lethal stress.

We did not address viable but nonculturable cells (VBNCs). Some studies indicate that VBNCs and persisters exhibit comparable phenotypes, with the primary distinction being that, following stress such as antibiotics, persisters can proliferate in both solid and liquid media. In contrast, VBNCs are said to only thrive in liquid media. There is an ongoing argument in the scientific community regarding whether VBNCs are genuinely persisters or merely dying cells or whether they exist[117-121]. Though it is often cited that the difference between VBNCs and persisters is that VBNCs do not grow on solid media, this defies the original definition of persisters proposed by Biggers, who discovered and named persisters[35]. Biggers described cultures that grow after lethal antibiotic treatment on agar plate as persisters, and also described cultures that grow in liquid after lethal antibiotic treatment as persisters. Though the original paper that identified VBNCs[122] did not mention persister, persisters were discovered first, nearly ∼40 years before VBNCs. This means anything that grows in liquid after lethal antibiotic treatment should be called persisters regardless of whether they grow in liquid or on solid media. Cells called VBNCs should be recognized as persisters or a subpopulation of persisters. Biggers did not distinguish between cultures that grow only on liquid from those that grow in both liquid and on solid plates, so recognizing VBNCs, if they do exist, as a subset of persisters is an important clarity that is often lacking in the literature.

Though previous work has claimed that VBNCs are almost at equal concentrations as that of CFU/mL[123, 124] (50% normal and 50% VBNCs), recent work has provided strong evidence that some of the high levels of reported VBNCs in log cultures may be a result of improper microscopy counts[125]. Regardless, we examined the potential revival of VBNCs through the ∼0.5 cell/well method. The ∼0.5 cell/well dilution technique is based on the CFU/mL of the bacterial culture as obtained on solid plates. If our dilution technique is correct, then a ∼0.5 cell/well dilution should yield growth in ∼48 wells out of 96 in a 96-well optical bottom plate. If VBNCs exist and multiply only in liquid media, then we should have ∼1 cell/well instead of ∼0.5 cell/well with growth in all 96 wells. Instead, we consistently got growth in ∼48 wells with 0.5 cell/well dilution, indicating non-interference of VBNCs. Our results were corroborated by growth in ∼24, ∼12, and ∼6 wells when we performed 0.25, 0.125, and 0.0625 cell/well dilutions, respectively. The R^2^ is 0.99 (Fig. S3C) when the number of wells with growth is plotted vs. cells/well based on CFU/mL. This technique may not have encompassed all the intricacies of VBNCs, and not every well is likely to originate from 1 cell. We previously identified primed cells when *E. coli* was diluted to ∼2 cells/well approach (based on CFU/ml) and treated with ampicillin[30]. So, we tested *B*. *megaterium*, and once again observed a significant variability with Cip treatment, a ∼60-fold difference (Fig. S3D). This indicates that even if VBNCs are present, they do not skew our results, and we still detect primed cells even when using ∼2 cells per well. Moreover, this underscores that the FT does not necessitate a 1-cell/well dilution.

Why do bacteria have both persisters and primed cells? As mentioned in the introduction, heterogeneity leads to reduced fitness in an optimal, steady-state environment. This is likely because (1) some cells are not growing as fast as they could otherwise, (2) some cells are not growing at all (persisters), or (3) some cells are using resources to prepare for potential threats (primed cells). One way this reduced fitness is minimized is by only having a small fraction of cells behaving differently; the smaller the number of cells not focused on growing and dividing as fast as possible, the lesser the adverse effects on the overall population’s fitness. Another way to reduce the burden is to have *heterogeneity* in the types of cells by having both persisters and primed cells. Because primed cells are still growing, it may reduce the fitness cost. In contrast, cells in the persister state have a greater fitness cost; every cell that is not dividing is not consuming at the optimal rate and is not reproducing.

There is still a lot we do not know about primed cells and persisters. We do not know which mechanisms control primed cell formation and how the epigenetic memory is passed on. We previously provided evidence that there is no single persister-gene that controls persister formation, and multiple pathways lead to persister formation[28]. It seems logical that the same is true for primed cells. We do not know if *B. megaterium* and *E. coli* primed cells form because of similar or different mechanisms. We do not know a lot about the switching between primed cells, persistence, and susceptible cells.

It seems unlikely that the change in gene expression that leads to the primed state is carried over to the persister state or after the cell emerges from persistence because the persister (non-dividing) state is so vastly different from the growing state of primed cells. We do not know if when cells emerge from persistence, they are equivalent to susceptible cells or transitioned to primed cells before fully becoming susceptible cells. Though we do not have evidence for the latter (persister cell < > primed cell < > susceptible cell), this seems like a logical intermediate step. We often describe cells emerging from persistence once the antibiotic stress is removed, but this description could be oversimplified. It seems unlikely cells have the capacity to detect when an antibiotic is gone. If cells had the ability to detect the presence of an antibiotic, then they should not die over time; they should simply stay dormant until the antibiotic is removed. Cells die over time in a growth media because of two possibilities: (1) they lack essential nutrients, leading to cell death, or (2) they exit the persister state trying to divide, leading to death by the antibiotics. There are ample nutrients in the media for the test we describe in this work, so the first reason seem unlikely. Most likely, cells that exit persistence are being killed because most antibiotics only kill cells that are dividing. The Woods group has proposed cells emerge from persistence when the hybridized ribosomes (trapped in a 100S state) are reactivated to 70S ribosomes[126]. This is an elegant way to describe the relatively rapid transition from persisters to a growing state. However, if some persisters emerge from persistence rapidly when the antibiotic level is high, they will be killed. A potential strategy to minimize the chance of being killed is to switch from persistence to a more stress-prepared state, and if the stress is still there, switch back to the persister state. At this point, we do not know how fast the switching between each state is.

We know that primed cells come from susceptible cells, and primed cells lead to more primed cells, susceptible cells, and persister cells (Fig. 1C). Previous work has provided evidence that extremely slow-growing cells can become persisters[78], but primed cells are not extremely slow-growing cells[30]. This contradiction suggests that not all persister cells come from primed cells. In addition, this work does not have the sensitivity to test if primed cells are growing slightly slower or at the same rate as susceptible cells. In addition, we do not know how much switching to the primed cell state affects the overall state of the cell.

Our results show that there are more persisters in the stationary phase but do not support the existence of Type I and Type II persisters (Type II persisters form in the log phase, while Type I persisters that form in the stationary phase)[31, 127]. Here, we will use a simple analogy to explain these results. Imagine that you travel the same distance to school each day. You can ride the school bus, and you can walk the same path as the bus. If you walk to school with an empty bookbag, it takes on average 45 minutes; however, if you walk with a full (heavy) bookbag, it takes longer, on average 60 minutes. In this case, the mean and median time travel may be different, but the CV value (standard deviation/mean) is likely to be quite similar because regardless of the heaviness of the bookbag, it is the same person walking. This is similar to what we see with primed cells. However, if you walk to school half of the days and ride the bus for the other half of the days, the outcome would be different. The bus should be faster, and its CV will be quite different from that of when you walk. Similarly, suppose Type I and Type II persisters are formed through different mechanisms. In that case, we should have seen different numbers for the CV, fold difference, and range (upper and lower limits of persisters) in our FT replicates. Our results do not support two unique forms of persisters called Type I and Type II. The greater number of persisters in post-log phases is due to more primed cells in the transition and stationary phases compared to the log phase.

It is extremely unlikely that we had stationary persisters carrying over in our FTs because we diluted the starting culture down to individual cells, and the clonal populations grew from those single cells over many hours. If we had stationary persisters carrying over in our FTs, they would likely have had slower growth. However, we have no evidence that slower growth shows a correlation with higher persister percentages (Fig. 4C).

Slowed cell growth is often used to indicate the presence of a cellular burden because it is one of the easiest aspects to measure. However, a cellular burden does not necessarily need to affect growth. Cell burdens could have other effects on cells such as increased ATP use, higher expression of specific genes, accumulation of intermediates, and many others. For example, fluorescent proteins such as GFP are often used as reporters because they are easily visualized and do not have an apparent effect on growth at medium-to-low levels of expression. However, these proteins consistently produce reactive oxygen species (ROS)[128-130]. Cells are able to adapt to the ROS (which causes network disruptions), so this burden does not significantly affect growth. Another example of a cellular burden that affects growth is the addition of sub-MIC antibiotics. MIC (minimum inhibitory concentration) is the lowest concentration of an antibiotic that has a negative effect on growth, but cells can still grow. At sub-MIC levels, antibiotics should have no apparent effect on growth, yet many studies have shown that even at these levels, there is a cellular burden, which cells adapt to, resulting in normal growth rates[131-133]. As cells transition from the log phase to the stationary phase, a burden can be detected by measuring growth rates. Other micro-burdens (e.g. ROS) could occur in the log phase in individual cells, but cells adapt to the burdens with little to no effect on the population growth rate. These micro-burdens cause network disruptions, which cells respond to, and this response (change in gene expression) could temporarily lead to cells being more prepared for deadly stresses (i.e. antibiotics). This prepared state (expression of specific genes) could be passed down several generations. This is what we think happens in primed cells. It is logical to predict that in the transition and stationary phases, there are more of these micro-burdens than in the log phase, which results in an overall increased level of primed cells.

## Materials & Methods

### Microbial strains, media, and buffer

*Bacillus megaterium* (basionym) de Bary ATCC 14581, also known as *Priestia megaterium,* was used in this work. The bacterial cultures were grown in Miller’s modification Lennox lysogeny broth (LB) liquid media or LB with agar. All cultures were incubated at 37° C and shaken at 250 or 300 rpm. Minimal Media B (MMB)[28, 134] was used to buffer dilutions.

### Growth curve

*B. megaterium* culture was grown overnight in LB media at 37° C and 250 rpm in a shaker incubator. The overnight culture was subcultured 1:100 and frozen at -80°C for dilution experiments or distributed evenly into the wells of a ThermoFisher Nunc MicroWell 96-Well Optical Bottom Plate with Polymer Base, with at least 3 wells containing only LB media acting as blank. The plate was incubated in a FLUOstar Omega microplate reader at 37 °C and 300 rpm for about 20h. Absorbance was measured at 600 nm at regular intervals. The blank was subtracted, and a semi-log graph of Log OD_600 nm_ vs Time was generated.

### Death curve

*B. megaterium* culture was grown overnight in LB media at 37° C and 250 rpm in a shaker incubator. The overnight culture was sub-cultured at 1:100 in three replicates. The replicates were grown until the log phase (OD_600_ ∼0.5), and a specific volume from each replicate was diluted and plated. This served as the CFU/mL for 0h. The remaining volume of culture in replicates was treated with Cip with a final concentration of 6.4 µg/mL and incubated at 37° C, 250 rpm. A certain volume of the cultures was taken out after 1 h, 2 h, 3 h, 4 h, 5 h, 12 h, and 24 h post-antibiotic addition, diluted, and plated. These served as the CFU/mL for the respective time points. Persister percentages were calculated for each replicate at each time point, and their logarithmic values were plotted against time to generate a semi-log death curve.

### Population-level persister percentage variation assay: Noise Control (NC)

A log phase (OD_600_ ∼0.5) *B. megaterium* culture, grown in LB media, having ∼1E+7 CFU/mL (taken directly or frozen at -80°C), was divided into 48 wells (to serve as 48 replicates) of a ThermoFisher Nunc MicroWell 96-Well Optical-Bottom Plate with Polymer Base. A specific volume of the cultures from each replicate was diluted and plated. The remaining volume in the plate was then treated with Cip with a final concentration of 6.4 µg/mL for 3 h, at 37°C and 300 rpm, in a FLUOstar Omega microplate reader. Post-treatment, cultures from each replicate were again diluted and plated. All the plates were incubated for 10-12 h at 37°C and scanned using a flatbed Epson Perfection V370 Photo scanner post-incubation. Custom scripts were run to count bacterial colonies from the scanned images. Persister percentages were calculated for each replicate by ratioing their CFUs/mL post-antibiotic treatment to CFUs/mL pre-antibiotic treatment. A similar procedure was followed while doing this assay at later phases of growth. *B. megaterium* cultures at the transient phase had an OD_600_ of ∼1.0 (∼2E+7 CFU/mL), and at the stationary phase had an OD_600 nm_ of ∼1.5 (∼4E+7 CFU/mL).

### Single cell-level persister percentage variation assay: Fluctuation Test (FT)

A log phase (OD_600_ ∼0.5) *B. megaterium* culture, grown in LB media (taken directly or frozen at -80°C), having ∼1E+7 CFU/mL, was diluted down to ∼0.5 cells per well (or 1 cell per 2 wells) in a ThermoFisher Nunc MicroWell 96-Well Optical Bottom Plate with Polymer Base and typically about 50% of the wells had growth (ideally 48, but typically 46-50 wells out of 96 wells). For every single cell-level persister percentage variation assay, the dilution procedure was done twice to confirm the accuracy. The plates were incubated in FLUOstar Omega microplate reader at 37° C, 300 rpm for 6-8 h, and the cultures were grown to log phase ∼1E+7 CFU/mL (∼OD_600_ 0.5). As per the dilution, each of the bacterial cultures in the 48 wells likely proliferated from a single cell within the respective wells. Thus, 48 wells serve as 48 clonal populations. A specific volume of the cultures from each clonal population was diluted and plated. The remaining volume of each clonal population in the plate was then treated with Cip with a final concentration of 6.4 µg/mL for 3 h, at 37° C and 300 rpm, in a FLUOstar Omega microplate reader. Post-treatment, cultures from each clonal population were again diluted and plated. Plates were incubated for 10-12 h at 37° C and scanned using a flatbed Epson Perfection V370 Photo scanner post-incubation. Custom scripts were run to count bacterial colonies from the scanned images. Persister percentages were calculated for each clonal population by ratioing their CFUs/mL post-antibiotic treatment to CFUs/mL pre-antibiotic treatment. A similar procedure was followed while doing this assay at later phases of growth.

### Antibiotic resistance assay

After each fluctuation test, multiple colonies from high-persister wells were picked and streaked with and without ciprofloxacin-containing LB agar plates and incubated at 37° C.

### Spore detection assay

Stationary phase cultures (OD_600_ ∼1.5) having ∼4E+7 CFU/mL were subjected to heat shock (100° C for 1 min) and plated to check for the presence of spores. No growth was detected on LB plates, which indicates no viable spore formation.

### Memory assay

A log phase (OD_600_ ∼0.5) *B. megaterium* culture having ∼1E+7 CFU/mL was diluted down to 0.5 cells per well as described above, and cultures were incubated in a microplate reader at 37° C, 300 rpm for 6-8 h, and the cultures were grown to log phase ∼1E+7 CFU/mL (∼OD_600_ 0.5). 48 wells serving 48 different clonal populations were selected for further experimentation. Then, they were diluted in prewarmed LB to 1:2, 1:10, and 1:100, respectively. Each dilution was performed in 2 replicates. Fluctuation Test (FT) was performed with the 2 replicates simultaneously for each dilution using Cip. The R^2^ values were determined from a regression line for the persister percentages of each well in both the replicates for all the dilutions.

### Fluoride assays

We chose sodium fluoride (NaF) as a non-antibiotic chemical stressor to test for primed cells. Both population-level and single cell-level persister percentage variation assays were performed in an identical way, as mentioned above, but with 60 mM NaF instead of Cip. MMB was used to buffer the reduction of pH by NaF in the culture media.

### Minimal inhibitory concentration (MIC) assay

A log phase (OD_600_ ∼0.5) *B. megaterium* culture, grown in LB media, was used to make a lawn on a Lennox L agar plate. A Liofilchem Cip MIC Test Strip[135] was then placed on the agar surface, and the plate was incubated at 37° C for 8-10h. The MIC value was determined as indicated by the tail of the zone of exclusion. Colonies from a high persister plate obtained through FT were re-cultured in LB media up to the log phase (OD_600_ ∼0.5) and were used to make a lawn on an LB/agar plate. MIC was determined similarly with a test strip. MIC of NaF was determined using a custom serial microdilution technique.

### Statistical analyses

Microsoft Excel, Spyder (Python version 3.12) and Rstudio programming applications were employed for post-hoc statistical data analysis and plotting. Statistical significance (two-tailed p-value) was determined using T-test with either similar or dissimilar variance (based on F-test results). The growth rate in Fig. 4C has been calculated using the formula:

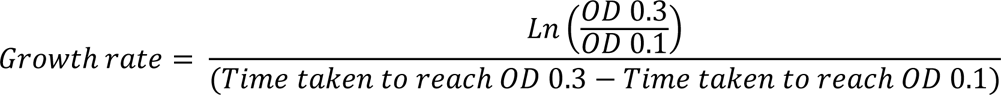

## Acknowledgements

We sincerely extend our gratitude to Hazera Khatun Koly for sharing her Cip MIC strip-test results with *B. megaterium*. This work is supported by:

- The National Science Foundation award numbers 1922542, 2240028, and 1849206,
- USDA National Institute of Food and Agriculture Hatch project grant number SD00H763-22, accession no. 7002192.
- The National Institute of Health (NIH-NIGMS) grant R35GM148351.

## Author contributions

MG performed the experiments, analyzed the data, and wrote the manuscript. AS and NCB conceived the project. NCB planned and directed it. All authors contributed to discussing and editing the manuscript.

## Data and materials availability

All data are available in the main text or the supplementary materials.

## Supplementary Materials

**Fig. S1.**
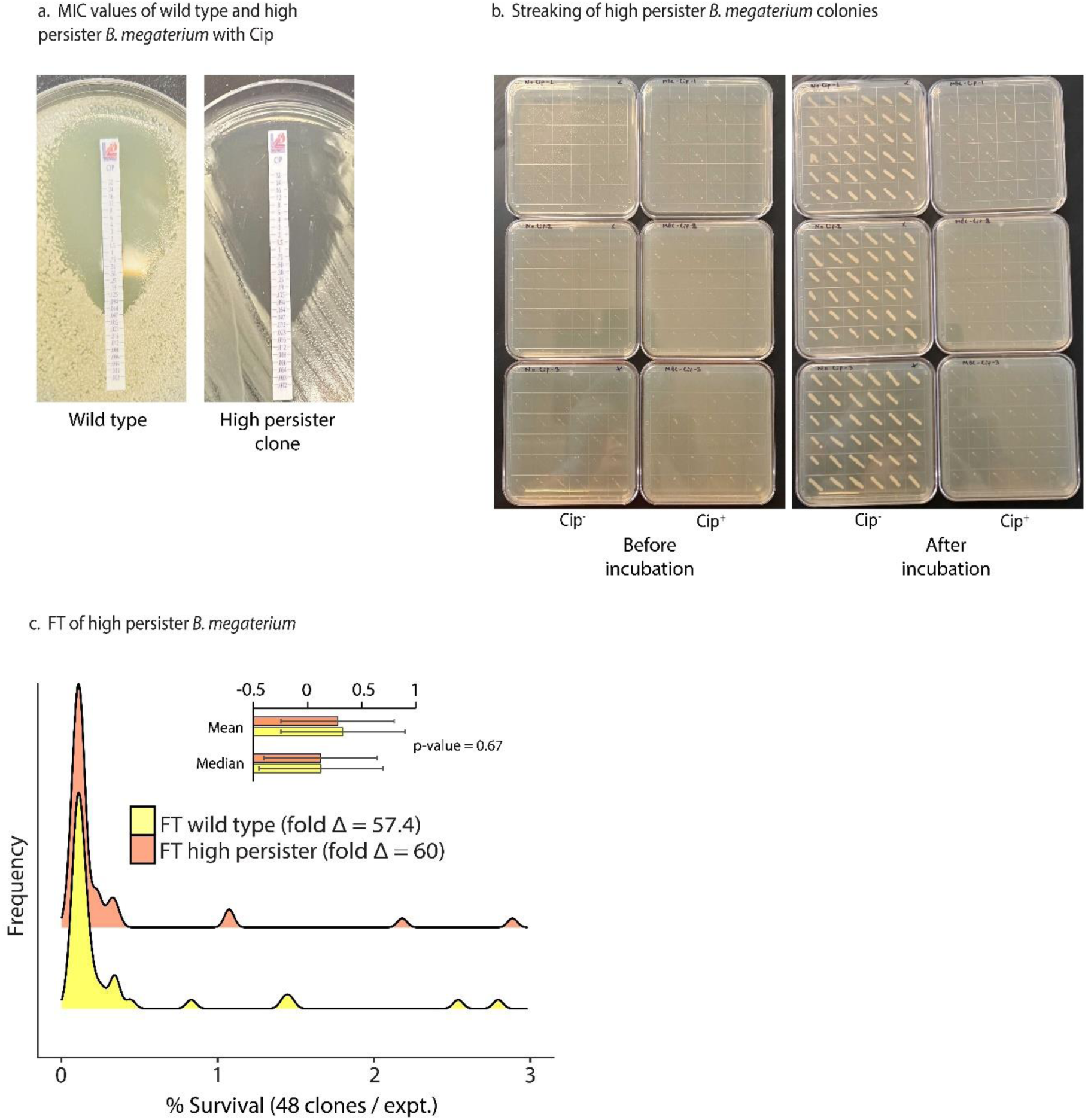
**(A).** MIC strip-test showing no change in MIC value of a high-persister clone. **(B).** The streaking of high-persister colonies shows that they are not resistant, eliminating mutation as the reason for the high variability in persister numbers. **(C).** Density plot of FT with a high-persister clone against Cip compared to WT. The high-persister clone yielded results similar to those of WT.

**Fig. S2.**
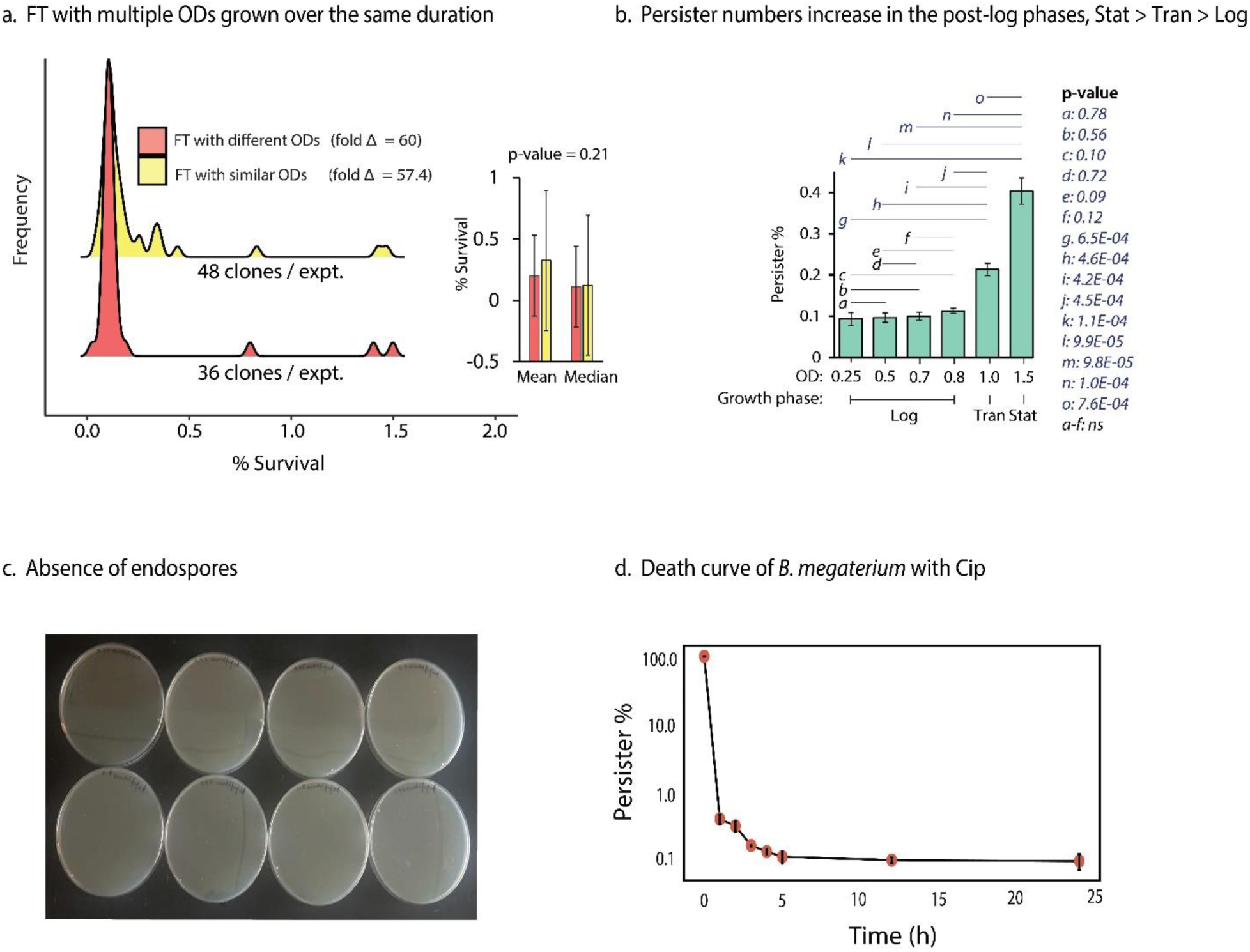
**(A).** FT with clones having different cell densities indicated through a wide range of ODs at 600 nm (0.25 – 0.8), grown over the same duration, yielded comparable results to that of wild-type. The average persister percentage and the range were similar. **(B).** Graph showing the persister percentages of *B. megaterium* against Cip at log phase (OD_600_ 0.25, 0.5, 0.7, 0.8), transition phase (OD_600_ 1.0) and stationary phase (OD_600_ 1.5). The persister percentages begin to increase post-log phase of growth. n = 3. The non-significant p-values are also indicated in this graph. **(C).** *B. megaterium* (at OD_600_ of 0.31, 0.43, 0.55. 0.7, 0.8, 1.02, 1.5 and 1.96) plated after 1min of heat shock at 100°C. No growth indicated a lack of endospores in the cultures. **(D).** Death curve of *B. megaterium* with lethal doses of Cip. The persister percentages have been measured through CFU/mL. The y-axis is in the log_10_ scale.

**Fig. S3.**
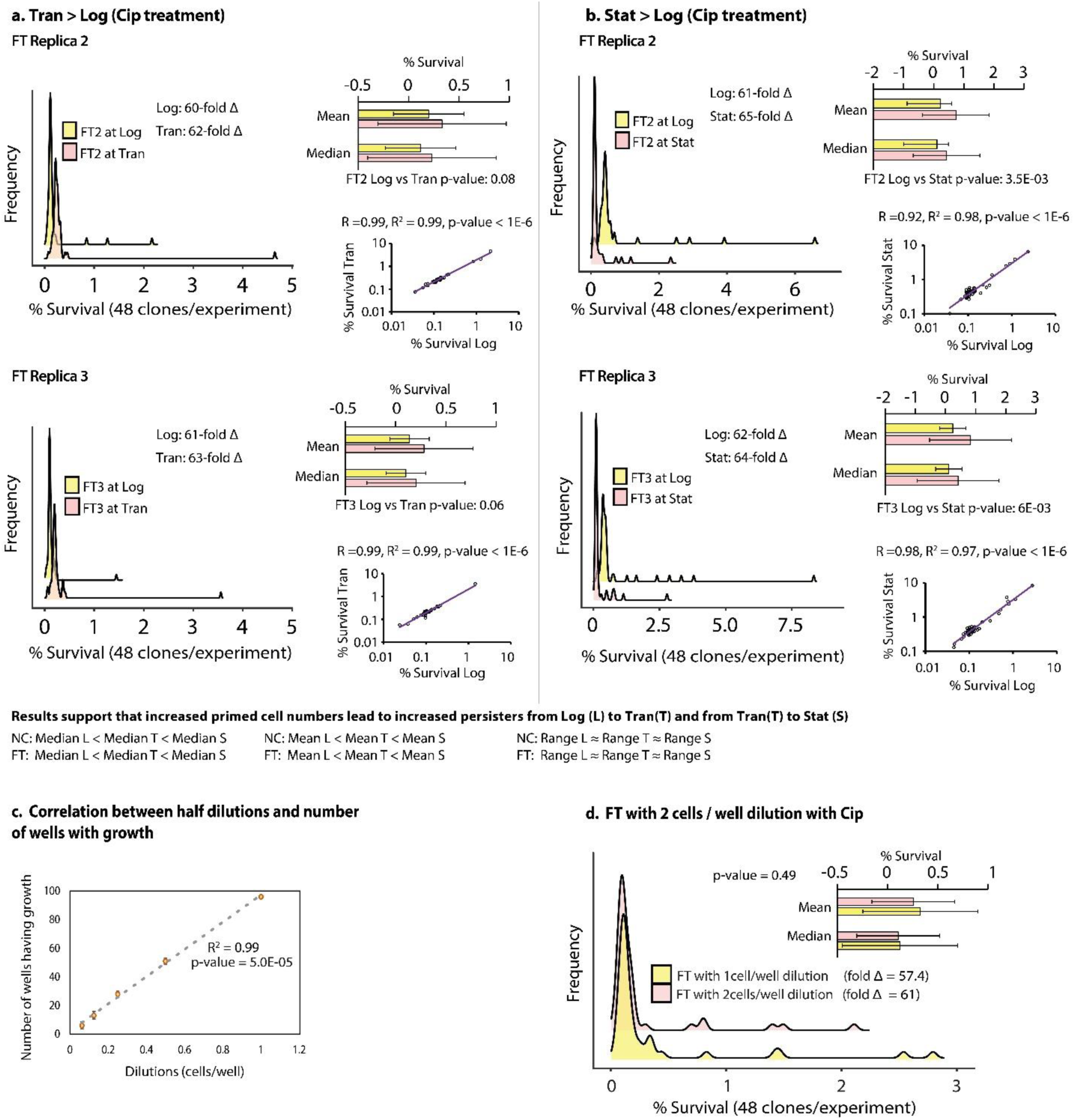
**(A).** Results of single cell-level persister percentage variation assay replicates (FT2 and FT3) at log (Log) vs transition (Tran), and **(B).** log (Log) vs stationary (Stat) phases of growth with Cip. For the correlation graphs, the x- and y-axes are on a log _10_ scale. **(C).** The graph shows the average number of wells that have growth with the dilutions (indicated by the x-axis values) performed per well. Total number of wells = 96. **(D).** Results of an FT against Cip performed with ∼2 cells per well yielded comparable results to that of one cell per well.

## References

1. Bhandari, M., et al., Genomic Diversity, Antimicrobial Resistance, Plasmidome, and Virulence Profiles of Salmonella Isolated from Small Specialty Crop Farms Revealed by Whole-Genome Sequencing. Antibiotics (Basel), 2023. 12(11).

2. Fenske, G.J. and J. Scaria, Analysis of 56,348 Genomes Identifies the Relationship between Antibiotic and Metal Resistance and the Spread of Multidrug-Resistant Non-Typhoidal Salmonella. Microorganisms, 2021. 9(7).

3. D’Costa, V.M., et al., Sampling the Antibiotic Resistome. Science, 2006. 311(5759): p. 374-377.

4. Giedraitienė, A., et al., Antibiotic Resistance Mechanisms of Clinically Important Bacteria. Medicina, 2011. 47(3): p. 19.

5. Cox, G. and G.D. Wright, Intrinsic antibiotic resistance: mechanisms, origins, challenges and solutions. International Journal of Medical Microbiology, 2013. 303(6-7): p. 287–292.

6. Murray, C.J., et al., Global burden of bacterial antimicrobial resistance in 2019: a systematic analysis. The Lancet, 2022. 399(10325): p. 629–655.

7. Huemer, M., et al., Antibiotic resistance and persistence—Implications for human health and treatment perspectives. EMBO reports, 2020. 21(12): p. e51034.

8. Control, C.f.D. and Prevention, Antibiotic resistance threats in the United States, 2019. 2019: US Department of Health and Human Services, Centres for Disease Control and ….

9. American Cancer Society. Global Cancer Facts & Figures in American Cancer Society. 2018.

10. ; 5th edition:[

11. Hall-Stoodley, L., J.W. Costerton, and P. Stoodley, Bacterial biofilms: from the natural environment to infectious diseases. Nature reviews microbiology, 2004. 2(2): p. 95–108.

12. Harms, A., E. Maisonneuve, and K. Gerdes, Mechanisms of bacterial persistence during stress and antibiotic exposure. Science, 2016. 354(6318): p. aaf4268.

13. Gollan, B., et al., Bacterial persisters and infection: past, present, and progressing. Annual review of microbiology, 2019. 73: p. 359–385.

14. Lewis, K., Persister Cells. Annual Review of Microbiology, 2010. 64(1): p. 357–372.

15. Brauner, A., et al., Distinguishing between resistance, tolerance and persistence to antibiotic treatment. Nature Reviews Microbiology, 2016. 14(5): p. 320–330.

16. Levin-Reisman, I., et al., Antibiotic tolerance facilitates the evolution of resistance. Science, 2017. 355(6327): p. 826-830.

17. Van den Bergh, B., et al., Frequency of antibiotic application drives rapid evolutionary adaptation of Escherichia coli persistence. Nature microbiology, 2016. 1(5): p. 1–7.

18. Windels, E.M., et al., Bacterial persistence promotes the evolution of antibiotic resistance by increasing survival and mutation rates. The ISME journal, 2019. 13(5): p. 1239–1251.

19. Long, A.M., et al., Benchmarking microbial growth rate predictions from metagenomes. The ISME Journal, 2021. 15(1): p. 183–195.

20. Kearns, D.B. and R. Losick, Cell population heterogeneity during growth of Bacillus subtilis. Genes Dev, 2005. 19(24): p. 3083–94.

21. Veening, J.-W., W.K. Smits, and O.P. Kuipers, Bistability, epigenetics, and bet-hedging in bacteria. Annu. Rev. Microbiol., 2008. 62: p. 193–210.

22. Kaern, M., et al., Stochasticity in gene expression: from theories to phenotypes. Nature Reviews Genetics, 2005. 6(6): p. 451–464.

23. Paulsson, J., Summing up the noise in gene networks. Nature, 2004. 427(6973): p. 415-418.

24. Raser, J.M. and E.K. O’shea, Noise in gene expression: origins, consequences, and control. Science, 2005. 309(5743): p. 2010-2013.

25. Symmons, O. and A. Raj, What’s Luck Got to Do with It: Single Cells, Multiple Fates, and Biological Nondeterminism. Mol Cell, 2016. 62(5): p. 788–802.

26. Engl, C., Noise in bacterial gene expression. Biochem Soc Trans, 2019. 47(1): p. 209–217.

27. Cohen, D., Optimizing reproduction in a randomly varying environment when a correlation may exist between the conditions at the time a choice has to be made and the subsequent outcome. Journal of Theoretical Biology, 1967. 16(1): p. 1–14.

28. Deter, H.S., T. Hossain, and N.C. Butzin, Antibiotic tolerance is associated with a broad and complex transcriptional response in E. coli. Scientific reports, 2021. 11(1): p. 6112.

29. Rahman, K.M.T., et al., An isogenic E. coli population gives rise to multiple persister phenotypes. bioRxiv, 2024. Submited to Journal: p. 2023.12.09.570944.

30. Hossain, T., A. Singh, and N.C. Butzin, Escherichia coli cells are primed for survival before lethal antibiotic stress. Microbiology Spectrum, 2023. 11(5): p. e01219-23.

31. Balaban, N.Q., et al., Bacterial persistence as a phenotypic switch. Science, 2004. 305(5690): p. 1622-5.

32. Keren, I., et al., Persister cells and tolerance to antimicrobials. FEMS microbiology letters, 2004. 230(1): p. 13–18.

33. Acar, M., J.T. Mettetal, and A. Van Oudenaarden, Stochastic switching as a survival strategy in fluctuating environments. Nature genetics, 2008. 40(4): p. 471–475.

34. Thattai, M. and A. Van Oudenaarden, Stochastic gene expression in fluctuating environments. Genetics, 2004. 167(1): p. 523–530.

35. Bigger, J., Treatment of Staphyloeoeeal Infections with Penicillin by Intermittent Sterilisation. Lancet, 1944: p. 497–500.

36. Raj, A. and A. Van Oudenaarden, Nature, nurture, or chance: stochastic gene expression and its consequences. Cell, 2008. 135(2): p. 216–226.

37. Brandt, L., S. Cristinelli, and A. Ciuffi, Single-cell analysis reveals heterogeneity of virus infection, pathogenicity, and host responses: HIV as a pioneering example. Annual review of virology, 2020. 7: p. 333–350.

38. Foreman, R. and R. Wollman, Mammalian gene expression variability is explained by underlying cell state. Molecular systems biology, 2020. 16(2): p. e9146.

39. Lyu, Z., et al., Heterogeneous flagellar expression in single salmonella cells promotes diversity in antibiotic tolerance. MBio, 2021. 12(5): p. 10.1128/mbio.02374-21.

40. SoRelle, E.D., et al., Single-cell RNA-seq reveals transcriptomic heterogeneity mediated by host– pathogen dynamics in lymphoblastoid cell lines. Elife, 2021. 10: p. e62586.

41. Van Eyndhoven, L.C., A. Singh, and J. Tel, Decoding the dynamics of multilayered stochastic antiviral IFN-I responses. Trends in immunology, 2021. 42(9): p. 824–839.

42. Topolewski, P., et al., Phenotypic variability, not noise, accounts for most of the cell-to-cell heterogeneity in IFN-γ and oncostatin M signaling responses. Science Signaling, 2022. 15(721): p. eabd9303.

43. Shaffer, S.M., et al., Memory sequencing reveals heritable single-cell gene expression programs associated with distinct cellular behaviors. Cell, 2020. 182(4): p. 947–959. e17.

44. Shaffer, S.M., et al., Rare cell variability and drug-induced reprogramming as a mode of cancer drug resistance. Nature, 2017. 546(7658): p. 431-435.

45. Chang, C.A., et al., Ontogeny and vulnerabilities of drug-tolerant persisters in HER2+ breast cancer. Cancer discovery, 2022. 12(4): p. 1022–1045.

46. Bokes, P. and A. Singh, A modified fluctuation test for elucidating drug resistance in microbial and cancer cells. European Journal of Control, 2021. 62: p. 130–135.

47. Singh, A. and M. Saint-Antoine, Probing transient memory of cellular states using single-cell lineages. Front Microbiol, 2022. 13: p. 1050516.

48. Luria, S.E. and M. Delbrück, Mutations of Bacteria from Virus Sensitivity to Virus Resistance. Genetics, 1943. 28(6): p. 491–511.

49. Saint-Antoine, M., R. Grima, and A. Singh, A fluctuation-based approach to infer kinetics and topology of cell-state switching. bioRxiv, 2022: p. 2022.03.30.486492.

50. Cohen, N.R., M.A. Lobritz, and J.J. Collins, Microbial persistence and the road to drug resistance. Cell Host Microbe, 2013. 13(6): p. 632–42.

51. Wallis, R.S., et al., Drug tolerance in Mycobacterium tuberculosis. Antimicrobial agents and chemotherapy, 1999. 43(11): p. 2600–2606.

52. Meylan, S., I.W. Andrews, and J.J. Collins, Targeting antibiotic tolerance, pathogen by pathogen. Cell, 2018. 172(6): p. 1228–1238.

53. Mulcahy, L.R., et al., Emergence of Pseudomonas aeruginosa strains producing high levels of persister cells in patients with cystic fibrosis. Journal of bacteriology, 2010. 192(23): p. 6191–6199.

54. LaFleur, M.D., Q. Qi, and K. Lewis, Patients with long-term oral carriage harbor high-persister mutants of Candida albicans. Antimicrobial agents and chemotherapy, 2010. 54(1): p. 39–44.

55. Schumacher, M.A., et al., HipBA–promoter structures reveal the basis of heritable multidrug tolerance. Nature, 2015. 524(7563): p. 59-64.

56. Van den Bergh, B., M. Fauvart, and J. Michiels, Formation, physiology, ecology, evolution and clinical importance of bacterial persisters. FEMS microbiology reviews, 2017. 41(3): p. 219–251.

57. Lewis, K., Persister cells, dormancy and infectious disease. Nature Reviews Microbiology, 2007. 5(1): p. 48–56.

58. Fisher, R.A., B. Gollan, and S. Helaine, Persistent bacterial infections and persister cells. Nature Reviews Microbiology, 2017. 15(8): p. 453–464.

59. Ye, M., et al., A modified limiting dilution method for monoclonal stable cell line selection using a real-time fluorescence imaging system: A practical workflow and advanced applications. Methods and Protocols, 2021. 4(1): p. 16.

60. Zitzmann, J., et al., Single-cell cloning enables the selection of more productive Drosophila melanogaster S2 cells for recombinant protein expression. Biotechnology Reports, 2018. 19: p. e00272.

61. Balaban, N.Q., et al., Definitions and guidelines for research on antibiotic persistence. Nature Reviews Microbiology, 2019. 17(7): p. 441–448.

62. Rafailidis, P.I., E.N. Ioannidou, and M.E. Falagas, Ampicillin/Sulbactam. Drugs, 2007. 67(13): p. 1829–1849.

63. O’Connor, S., et al., Apramycin, a unique aminocyclitol antibiotic. J Org Chem, 1976. 41(12): p. 2087–92.

64. LeBel, M., Ciprofloxacin: chemistry, mechanism of action, resistance, antimicrobial spectrum, pharmacokinetics, clinical trials, and adverse reactions. Pharmacotherapy, 1988. 8(1): p. 3–33.

65. Jablonka, E. and M.J. Lamb, Evolution in four dimensions, revised edition: Genetic, epigenetic, behavioral, and symbolic variation in the history of life. 2014: MIT press.

66. Casadesús, J. and D. Low, Epigenetic gene regulation in the bacterial world. Microbiology and molecular biology reviews, 2006. 70(3): p. 830–856.

67. Casadesús, J. and R. D’Ari, Memory in bacteria and phage. Bioessays, 2002. 24(6): p. 512–518.

68. Lisman, J.E., A mechanism for memory storage insensitive to molecular turnover: a bistable autophosphorylating kinase. Proceedings of the National Academy of Sciences, 1985. 82(9): p. 3055–3057.

69. Rando, O.J. and K.J. Verstrepen, Timescales of genetic and epigenetic inheritance. Cell, 2007. 128(4): p. 655–668.

70. Libby, E.A., S. Reuveni, and J. Dworkin, Multisite phosphorylation drives phenotypic variation in (p)ppGpp synthetase-dependent antibiotic tolerance. Nature Communications, 2019. 10(1): p. 5133.

71. Xu, Y., et al., DNA adenine methylation is involved in persister formation in E. coli. Microbiol Res, 2021. 246: p. 126709.

72. Murawski, A.M. and M.P. Brynildsen, Ploidy is an important determinant of fluoroquinolone persister survival. Current Biology, 2021. 31(10): p. 2039–2050. e7.

73. Harsh, V., K. Maryam, and S. Hanna, Non-genetic inheritance restraint of cell-to-cell variation. eLife, 2021. 10.

74. Lewis, K., Persister cells. Annual review of microbiology, 2010. 64: p. 357–372.

75. Etienne, M. and G. Kenn, Molecular Mechanisms Underlying Bacterial Persisters. Cell, 2014. 157(3): p. 539–548.

76. Maisonneuve, E. and K. Gerdes, Molecular mechanisms underlying bacterial persisters. Cell, 2014. 157(3): p. 539–548.

77. Möker, N., C.R. Dean, and J. Tao, Pseudomonas aeruginosa increases formation of multidrug-tolerant persister cells in response to quorum-sensing signaling molecules. Journal of bacteriology, 2010. 192(7): p. 1946–1955.

78. Kussell, E., et al., Bacterial persistence: a model of survival in changing environments. Genetics, 2005. 169(4): p. 1807–14.

79. Hengge-Aronis, R., Signal transduction and regulatory mechanisms involved in control of the sigma(S) (RpoS) subunit of RNA polymerase. Microbiol Mol Biol Rev, 2002. 66(3): p. 373–95, table of contents.

80. Wood, T.K., S.J. Knabel, and B.W. Kwan, Bacterial persister cell formation and dormancy. Appl Environ Microbiol, 2013. 79(23): p. 7116–21.

81. Lopatkin, A.J., et al., Bacterial metabolic state more accurately predicts antibiotic lethality than growth rate. Nature Microbiology, 2019. 4(12): p. 2109–2117.

82. Cañas-Duarte, S.J., S. Restrepo, and J.M. Pedraza, Novel protocol for persister cells isolation. PLoS One, 2014. 9(2): p. e88660.

83. Petersen, P.E. and M.A. Lennon, Effective use of fluorides for the prevention of dental caries in the 21st century: the WHO approach. Community dentistry and oral epidemiology, 2004. 32(5): p. 319–321.

84. Arnold Jr, F., et al., Fifteenth year of the Grand Rapids fluoridation study. The Journal of the American Dental Association, 1962. 65(6): p. 780–785.

85. Blayney, J.R. and I.N. Hill, Fluorine and dental caries. Journal of the American Dental Association (1939), 1967. 74(2): p. 225-302.

86. Hutton, W.L., B.W. Linscott, and D.B. Williams, Final report of local studies on water fluoridation in Brantford. Canadian Journal of Public Health/Revue Canadienne de Sante’e Publique, 1956. 47(3): p. 89–92.

87. Ast, D.B. and B. Fitzgerald, Effectiveness of water fluoridation. The Journal of the American Dental Association, 1962. 65(5): p. 581–587.

88. Marquis, R., Diminished acid tolerance of plaque bacteria caused by fluoride. Journal of Dental Research, 1990. 69(2_suppl): p. 672-675.

89. Marquis, R.E., Antimicrobial actions of fluoride for oral bacteria. Can J Microbiol, 1995. 41(11): p. 955–64.

90. Pedersen, J.T., et al., Metal fluoride inhibition of a P-type H+ pump: stabilization of the phosphoenzyme intermediate contributes to post-translational pump activation. Journal of Biological Chemistry, 2015. 290(33): p. 20396–20406.

91. Sturr, M.G. and R.E. Marquis, Inhibition of proton-translocating ATPases of Streptococcus mutans and Lactobacillus casei by fluoride and aluminum. Archives of microbiology, 1990. 155: p. 22–27.

92. Bunick, F.J. and S. Kashket, Enolases from fluoride-sensitive and fluoride-resistant streptococci. Infection and Immunity, 1981. 34(3): p. 856–863.

93. Song, C., et al., Sodium fluoride induces nephrotoxicity via oxidative stress-regulated mitochondrial SIRT3 signaling pathway. Scientific Reports, 2017. 7(1): p. 672.

94. Thibodeau, E.A. and T.F. Keefe, pH-dependent fluoride inhibition of catalase activity. Oral microbiology and immunology, 1990. 5(6): p. 328–331.

95. Forbes, S., et al., Simultaneous assessment of acidogenesis-mitigation and specific bacterial growth-inhibition by dentifrices. PloS One, 2016. 11(2): p. e0149390.

96. Guha-Chowdhury, N., Y. Iwami, and T. Yamada, Effect of low levels of fluoride on proton excretion and intracellular pH in glycolysing streptococcal cells under strictly anaerobic conditions. Caries research, 1997. 31(5): p. 373–378.

97. Bowen, W. and M.J. Hewitt, Effect of fluoride on extracellular polysaccharide production by Streptococcus mutans. Journal of Dental Research, 1974. 53(3): p. 627–629.

98. Ma, H., et al., Effects of fluoride on bacterial growth and its gene/protein expression. Chemosphere, 2014. 100: p. 190–193.

99. Wegman, M., et al., Effects of fluoride, lithium, and strontium on intracellular polysaccharide accumulation in S. mutans and A. viscosus. Journal of Dental Research, 1984. 63(9): p. 1126–1129.

100. Elowitz, M.B., et al., Stochastic gene expression in a single cell. Science, 2002. 297(5584): p. 1183-1186.

101. Sampaio, N.M.V. and M.J. Dunlop, Functional roles of microbial cell-to-cell heterogeneity and emerging technologies for analysis and control. Current opinion in microbiology, 2020. 57: p. 87–94.

102. Süel, G.M., et al., An excitable gene regulatory circuit induces transient cellular differentiation. Nature, 2006. 440(7083): p. 545-550.

103. Maamar, H., A. Raj, and D. Dubnau, Noise in gene expression determines cell fate in Bacillus subtilis. Science, 2007. 317(5837): p. 526-529.

104. Eldar, A. and M.B. Elowitz, Functional roles for noise in genetic circuits. Nature, 2010. 467(7312): p. 167-173.

105. Chalancon, G., et al., Interplay between gene expression noise and regulatory network architecture. Trends in genetics, 2012. 28(5): p. 221–232.

106. Johnston, I.G., et al., Mitochondrial variability as a source of extrinsic cellular noise. PLoS computational biology, 2012. 8(3): p. e1002416.

107. Neuert, G., et al., Systematic identification of signal-activated stochastic gene regulation. Science, 2013. 339(6119): p. 584-587.

108. Dar, R.D., et al., Screening for noise in gene expression identifies drug synergies. Science, 2014. 344(6190): p. 1392-1396.

109. Magklara, A. and S. Lomvardas, Stochastic gene expression in mammals: lessons from olfaction. Trends in cell biology, 2013. 23(9): p. 449–456.

110. Battich, N., T. Stoeger, and L. Pelkmans, Control of transcript variability in single mammalian cells. Cell, 2015. 163(7): p. 1596–1610.

111. Larsson, A.J., et al., Genomic encoding of transcriptional burst kinetics. Nature, 2019. 565(7738): p. 251-254.

112. Singh, A., et al., Transcriptional bursting from the HIV-1 promoter is a significant source of stochastic noise in HIV-1 gene expression. Biophysical journal, 2010. 98(8): p. L32–L34.

113. Rodriguez, J., et al., Intrinsic dynamics of a human gene reveal the basis of expression heterogeneity. Cell, 2019. 176(1): p. 213–226. e18.

114. Larsson, A.J., et al., Transcriptional bursts explain autosomal random monoallelic expression and affect allelic imbalance. PLoS computational biology, 2021. 17(3): p. e1008772.

115. Fraser, L.C., et al., Reduction in gene expression noise by targeted increase in accessibility at gene loci. Proceedings of the National Academy of Sciences, 2021. 118(42): p. e2018640118.

116. Ochiai, H., et al., Genome-wide kinetic properties of transcriptional bursting in mouse embryonic stem cells. Science advances, 2020. 6(25): p. eaaz6699.

117. Song, S. and T.K. Wood, ‘Viable but non-culturable cells’ are dead. Environmental microbiology, 2021. 23(5): p. 2335–2338.

118. Kirschner, A.K., et al., How dead is dead? Viable but non-culturable versus persister cells. Environmental Microbiology Reports, 2021. 13(3): p. 243–245.

119. Nyström, T., Nonculturable bacteria: programmed survival forms or cells at death’s door? Bioessays, 2003. 25(3): p. 204–211.

120. McDougald, D., et al., Nonculturability: adaptation or debilitation? FEMS microbiology ecology, 1998. 25(1): p. 1–9.

121. Nyström, T., Not quite dead enough: on bacterial life, culturability, senescence, and death. Archives of microbiology, 2001. 176: p. 159–164.

122. Xu, H.S., et al., Survival and viability of nonculturableEscherichia coli andVibrio cholerae in the estuarine and marine environment. Microb Ecol, 1982. 8(4): p. 313–23.

123. Orman, M.A. and M.P. Brynildsen, Establishment of a method to rapidly assay bacterial persister metabolism. Antimicrobial agents and chemotherapy, 2013. 57(9): p. 4398–4409.

124. Ayrapetyan, M., T. Williams, and J.D. Oliver, Relationship between the viable but nonculturable state and antibiotic persister cells. Journal of bacteriology, 2018. 200(20): p. 10.1128/jb.00249-18.

125. Rahman, K.M.T. and N.C. Butzin, Counter-on-chip for bacterial cell quantification, growth, and live-dead estimations. Scientific Reports, 2024. 14(1).

126. Song, S. and T.K. Wood, ppGpp ribosome dimerization model for bacterial persister formation and resuscitation. Biochem Biophys Res Commun, 2020. 523(2): p. 281–286.

127. Aedo, S.J., M.A. Orman, and M.P. Brynildsen, Stationary phase persister formation in Escherichia coli can be suppressed by piperacillin and PBP3 inhibition. BMC Microbiol, 2019. 19(1): p. 140.

128. Dixit, R. and R. Cyr, Cell damage and reactive oxygen species production induced by fluorescence microscopy: effect on mitosis and guidelines for non-invasive fluorescence microscopy. Plant J, 2003. 36(2): p. 280–90.

129. Kalyanaraman, B. and J. Zielonka, Green fluorescent proteins induce oxidative stress in cells: A worrisome new wrinkle in the application of the GFP reporter system to biological systems? Redox Biol, 2017. 12: p. 755–757.

130. Ganini, D., et al., Fluorescent proteins such as eGFP lead to catalytic oxidative stress in cells. Redox Biol, 2017. 12: p. 462–468.

131. Kwon, Y.W. and S.Y. Lee, Effects of antibiotics at sub-minimal inhibitory concentrations on the morphology of Streptococcus mutans and Lactobacillus acidophilus. Oral Biology Research, 2020. 44(1): p. 1–7.

132. Chadha, J., In vitro effects of sub-inhibitory concentrations of amoxicillin on physiological responses and virulence determinants in a commensal strain of Escherichia coli. J Appl Microbiol, 2021. 131(2): p. 682–694.

133. Dong, G., et al., Effects of sub-minimum inhibitory concentrations of ciprofloxacin on biofilm formation and virulence factors of Escherichia coli. Braz J Infect Dis, 2019. 23(1): p. 15–21.

134. Deter, H.S., et al., Proteolytic queues at ClpXP increase antibiotic tolerance. ACS synthetic biology, 2019. 9(1): p. 95–103.

135. Wayne, P., Clinical and Laboratory Standards Institute: Performance standards for antimicrobial susceptibility testing: 20th informational supplement. CLSI document M100–S20, 2010.

